# Scorpion navigation by chemo-textural familiarity: modeling the interplay between sensory and landscape parameters

**DOI:** 10.1101/2025.08.18.670885

**Authors:** Douglas D Gaffin, Mariëlle H Hoefnagels

**Affiliations:** School of Biological Sciences, University of Oklahoma, Norman, Oklahoma, USA

## Abstract

The ability to return consistently to one point – e.g., a food source or shelter – is a poorly understood animal behavior. One hypothesis proposed to explain navigation in bees and ants is navigation by scene familiarity. In this paper, we propose that scorpions use ground-directed sensors (instead of vision) to augment navigation. Specifically, the paired pectines may detect matrices of chemical and textural information and compare it to stored memories, allowing the animal to step toward the most familiar “scene.” We first show that this form of familiarity navigation is consistent with scorpion homing excursions. We then developed a familiarity-based computer-simulated agent and analyzed navigation success using multiple combinations of sensor sensitivity levels, sensor resolutions, and landscape characteristics. We also assessed the effects of additional sensory channels and of landscape disruption. Finally, we showed that the simulated agent can navigate across a landscape the size of a typical sand scorpion’s home range.

## Introduction

Navigation by animals, especially those lacking obvious cognitive capacity, has enchanted and bewildered scientists for centuries. For example, how a honeybee with a brain size less than a cubic millimeter [1] navigates kilometers between a food source and its hive is a contentious debate [2–5]. Proposed mechanisms for hymenopteran navigation include a cognitive map of the animal’s environment [6, 7], landmark guidance [8], path integration [9], and navigation by scene familiarity [10, 11], sometimes termed view-based navigation [12, 13]. Of all these hypotheses, navigation by scene familiarity appears most parsimonious and has been implemented effectively in robot guidance [14].

Navigation by scene familiarity requires a sensor of adequate complexity and capacity to detect intricate patterns in the environment (e.g., the dense pixellated arrays of light-detecting ommatidia in the hymenopteran eye). It also requires a way to load into memory a collection of home-directed scenes acquired during initial short forays away from home (e.g., the learning flights of bees). With those ingredients in place, the rules of familiarity navigation are simple. An ant or bee that is returning to its nest or hive looks around itself and compares each scene to home-directed scenes previously stored in its memory. The animal then moves in the direction of the closest match and repeats the comparison for its next “step” [10, 15–17]. In other words, the insect navigates back to its home by choosing a direction that maximizes “familiarity.”

Sand scorpions offer an interesting alternative for investigating familiarity navigation while also extending the idea to sensory modalities other than vision. Sand scorpions are faithful to burrows they dig in the sand [18]. They make night-time hunting forays away from their burrows and return via inbound paths that do not retrace their outbound paths [19–22]. No one knows how they relocate their burrow, but accumulating evidence suggests they do it by following the same familiarity rules as hymenopterans [20, 23, 24], but instead of using light-detecting ommatidia, they use arrays of chemo-tactile sensilla arranged on ground-directed, midventral organs called pectines.

Much of the existing knowledge about pectines and scorpion behavior aligns with the requirements of familiarity navigation. Behavioral [25–30], morphological [31–37], and physiological [38–41] evidence indicates that the pectines are organs of taste and touch [42]. The pectines have multiple teeth that support dense matrices of peg sensilla (“pegs”), which detect mechanical [31, 38, 43] and near-range chemical [38, 40, 41] stimuli Fig 1. The pegs, which in some species exceed 100,000 in number, are functionally redundant [26, 44], suggesting sensory capability in line with the visual complexity of ommatidia in compound eyes [45]. The pectinal teeth are topographically mapped to the CNS, suggesting that spatial information from the pectines is preserved in the scorpion brain [34, 35, 46–49]. An unusual plexus of synapses among the peg sensilla [33, 39, 50, 51] appears to limit sensory adaptation, so the incoming matrix information remains “fresh” as the animal moves [52]. Finally, both path integration [21] and learning walks [53] in scorpions are potential mechanisms for gathering the home-directed chemical and/or textural information required for subsequent navigation by familiarity [9, 20, 24, 54, 55].peg sensi So, the ingredients are in place to suggest that scorpions are navigating by chemo-textural familiarity and that the pectines are the crucial organs of transduction. We had two main objectives for this study: 1) To assess if scorpion homing behavior is consistent with familiarity navigation. 2) To thoroughly test a computer model of pecten-based navigation, termed *Navigation by Familiarity with a Local Sensor* (NFLS) [56]. In the computer simulation, we tested navigation performance under various combinations of sensor parameters (resolution and sensitivity) and amount of landscape blur. We also tested the effects of displacement from the navigation path, adding sensory channels, and disrupting the landscape. Overall, this study shows that familiarity navigation using a downward-facing system of matrix-collecting sensors is plausible for scorpions.

**Fig 1.**
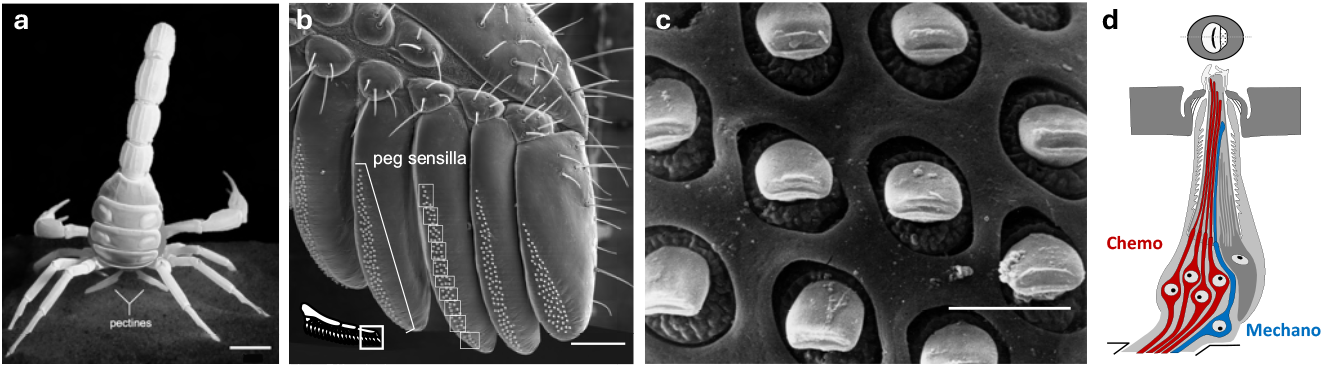
Scorpion pectines. (a) Posterior ventral view of female *Hadrurus arizonensis*, with the paired pectines brushing the ground. (b) SEM of the distal five teeth on the right pecten of a female *Paruroctonus utahensis* reveals dense matrices of minute peg sensilla. The superimposed rectangles each contain approximately eight sensilla, giving a sense of the putative chemosensory resolution of the tooth matrices [41]. (c) SEM of an expanded patch of peg sensilla from female *Smeringurus mesaensis* shows the consistent orientation of the slit-shaped terminal pores. (d) Simplified drawing of a longitudinal cut through a peg sensillum, depicting a population of chemosensory neurons (red) that extend into the peg shaft and a single mechanosensory neuron (blue) that terminates near the peg base. Photo credits: M. Hoefnagels (a), E. Knowlton (b), P. Brownell (c). [Scale bars: a = 1 cm, b = 50 µm, c = 5 µm]

## Results

### Assessment of home-directed journeys

In our previous laboratory investigation of *P. utahensis* homing behavior, we found that each animal dug an initial burrow in a small sand mound in the center of a larger circular arena, after which it made looping walks away from and back to the point of digging [53]. An example of these putative learning walks is illustrated in Fig 2; several additional examples are shown in S1 Fig. Note what the animal did on longer excursions after it completed the looping walks: On inbound journeys to the burrow region, the animal tended to select a consistent, relatively straight path (“directed movements” in Fig 2b,c). Furthermore, the outbound paths did not show signs of similarly directed behavior. This observation is consistent with the idea that the scorpion used information collected during learning walks to guide its way back to the burrow using navigation by familiarity. (S2 Fig provides an in-depth analysis of the same animal’s movements throughout the recording period.)

**Fig 2.**
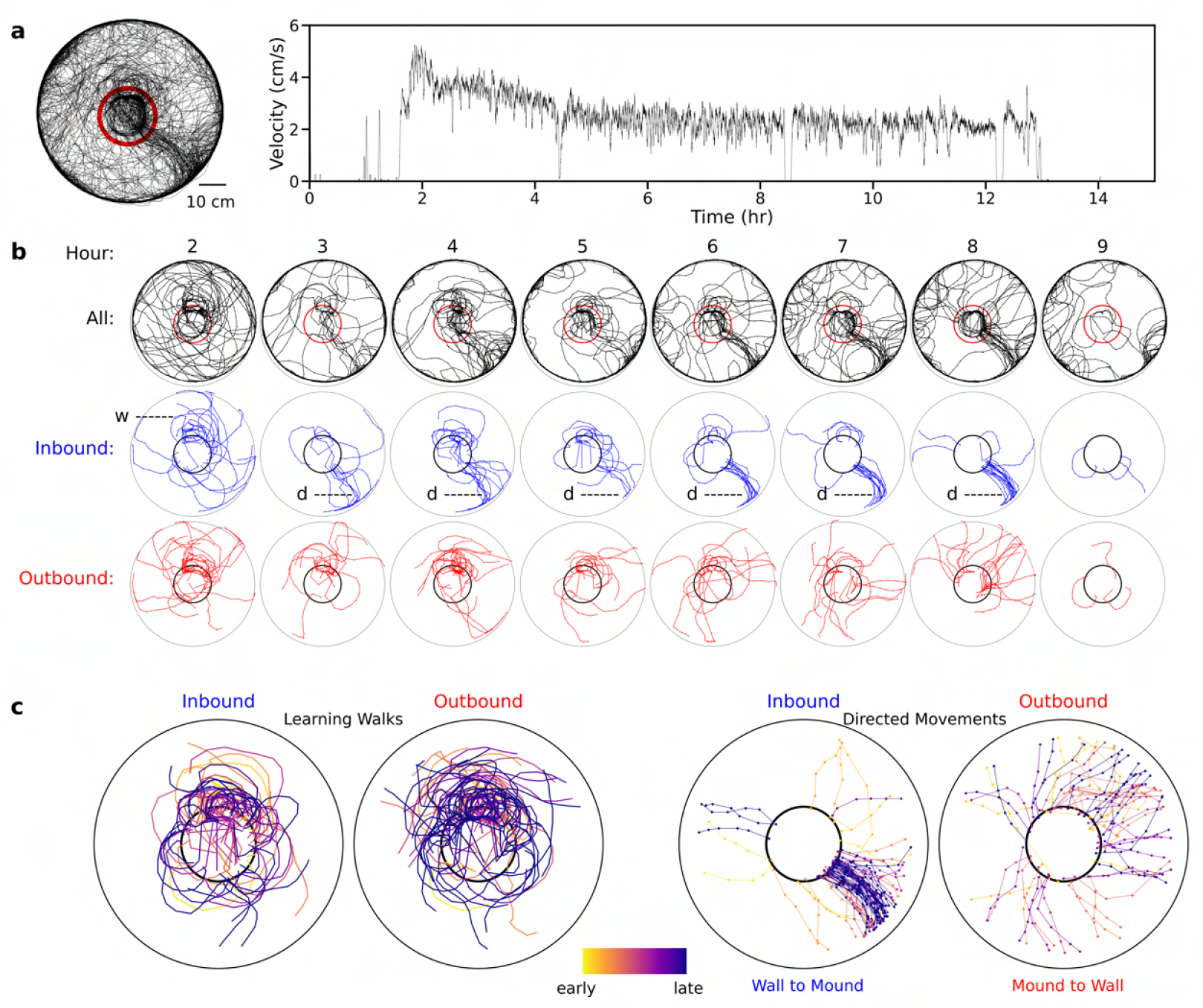
Learning walks and directed movements are consistent with familiarity navigation. (a) An all-night plot (15h, plotted at 2s intervals) of the movements of a sand scorpion (*P. utahensis*) in a sand-lined circular arena equipped with a small central mound of sand in which the animal dug a home burrow. The red circle outlines the burrow region used to extract inbound and outbound paths. The graph at right shows average velocity (cm/s; 60s moving average applied) of the animal for 15 hours, lasting from initial monitoring (5:00 PM) until after the animal returned to its burrow the following morning (7:00 AM). (b) The animal’s movements are parsed for hours 2-9 with all movements plotted at top in black. The “Inbound” paths (blue) show the 20 seconds of animal movement prior to crossing into the encircled area and the “Outbound” paths (red) show the subsequent 20 seconds of animal movement after exiting the encircled area. Putative learning walks (“w”) are indicated for the inbound plots during hour 2 and directed homeward movements (“d”) are indicated for hours 3-8. (c) The two plots at left show inbound and outbound paths that did not extend to within a few centimeters of the arena wall (putative learning walks). The two plots at right show full wall-to-mound and mound-to-wall plots. The paths are color-coded from “early” to “late” based on the key.

### Modeling familiarity navigation with a local sensor

Scorpions have pectines with the sensory complexity needed for navigation by chemo-textural familiarity, and their homing behavior is also consistent with this possibility. To further investigate this possibility, we developed a computer model to simulate pecten-based navigation. Our NFLS model digitally simulates the homing behavior observed above and in other behavioral accounts [18, 19, 22, 53, 57] by acquiring home-directed sensory information on a simulated landscape and using it to inform navigation during subsequent inbound journeys. We adapted our model from previous efforts [16, 17, 23, 56] and adjusted landscape blur, sensor resolution, and sensor sensitivity to determine the optimal combination of characteristics required for successful navigation.

### Acquisition of home-directed glimpses – the IJ path

The first step in modeling the navigation process is to create a path representing the *Initial Journey* (IJ) to simulate those generated by path integration and/or learning walks [20, 21, 53]. The path begins at an arbitrarily selected point in a simulated landscape and ends at the navigational goal (e.g., home burrow). We also created simulated pectines that acquire tens to thousands (depending on IJ path length) of “glimpses” of the substrate (at one glimpse per step) along the IJ path. As they are stored into memory, the glimpses are transformed based on the current user-set sensor resolution and number of sensor sensitivity levels (see variables in Methods). Although “glimpse” technically refers to visual stimuli, we use the term “glimpse” to refer to brief intermittent touches and/or tastes of the substrate via the peg sensilla matrices. In addition, we use the term “agent” to refer to the navigating scorpion on which the simulated pectines reside.

### Navigating by familiarity – the SJ path

To begin the *Subsequent Journey* (SJ), the agent is first placed near the beginning of the IJ path. The agent then spins 360° at 1° increments, comparing each current glimpse to all IJ glimpses stored in memory (e.g., if the agent collected 100 glimpses along the 100-step IJ, then 36000 comparisons are made). More specifically, the absolute differences on a pixel-by-pixel basis of each of the 360 current glimpses are subtracted from each IJ path glimpse and summed to produce an SAD value. SAD stands for the **Sum** of the **Absolute** pixel-by-pixel **Differences** of the matrices being compared (21). (The smaller the SAD value between any two glimpses, the more similar the two glimpses are.) The lowest SAD value is identified. The agent then checks which angle is associated with that value and steps forward a predefined distance in that direction. For each subsequent step, the agent rotates in a predefined, user-set arc (saccade), with each 1° glimpse again being compared to the entire IJ memory set. As before, the angle associated with the minimum SAD is selected, and the agent moves a predefined step length in that direction. This act of comparing rotational glimpses to those stored in memory is termed a *Rotational Information Difference Function* (RIDF) [12, 17, 58]. A related process, a *Translational Information Difference Function* (TIDF), [12, 58] is similar but instead of comparing pectinal glimpses of the substrate at a given point, the agent assesses glimpses displaced laterally left and right from its main heading. In this study we have applied RIDF techniques both separately and in concert with TIDF techniques. The above routines are iterated until the agent either gets within a defined distance of its goal (home) or goes too far astray.

A simplified version of the NFLS process is depicted in Fig 3, and a video showing the actual navigation of the agent is shown in S1 Video. We encourage readers to view the video, as it shows the incremental steps of the NFLS process that cannot be portrayed in Fig 3. The variables used to test the NFLS model are described in Methods.

**Fig 3.**
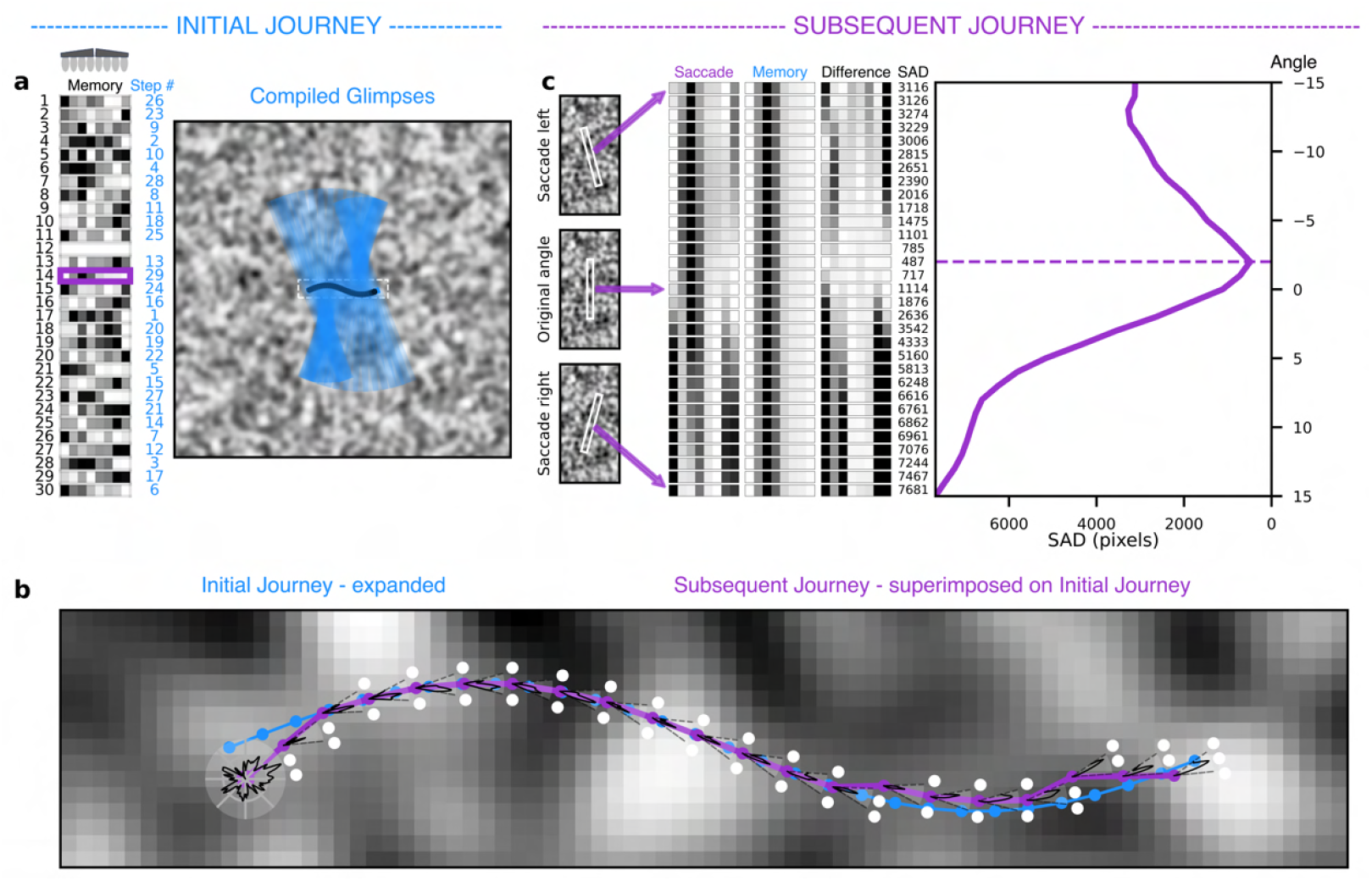
Simplified example of navigation by familiarity model. (a) Pectines are greatly simplified to 8 teeth (4 right, 4 left) and each tooth represents a unit of information. A short segment of an IJ is superimposed atop a small (1.16 x 1.16 cm) simulated landscape (grey levels = 255; Gaussian blur = 3.0). Here, 29 pectinal glimpses (one 8-tooth glimpse per step) were acquired as indicated by the compiled blue rectangles. The glimpses randomly fill 29 of the 30 memory cells at left (blue numbers at right indicate IJ step number). (b) To initiate an SJ (purple line), the agent is first placed near the beginning of the IJ (blue line). The agent rotates 360° and at each 1° turn computes the minimum SAD of the matrices being compared relative to the 29 stored glimpses. The polar plot (RIDF) in the lower left of (b) shows the summary of these computations (radial axis inverted). The agent then steps in the direction of the smallest SAD value and repeats the RIDF at this new point with its saccade restricted to a user-set angle to the left and right of its forward-facing direction (in this case +15° for a total of 30 saccadic glimpses). The agent gains translational information (TIDF) by repeating this RIDF at a set distance left and right of the original center position and steps forward from the point with the lowest SAD. The RIDF calculations for one step are visualized in (c) with the stack of 30 saccadic glimpses at left, a stack of 30 replications of an interrogated IJ glimpse from memory in the middle, and a stack of the absolute tooth-by-tooth differences at right, followed by a column showing the SAD values. This current saccade graph is the right most (final SJ step) wedge-shaped polar plot in (b) and the selected memory cell with the lowest SAD is highlighted by a purple outline in (a).

### Navigation success vs landscape and sensor variables

The navigation success of our NFLS model depended on both landscape and sensor parameters. For example, an agent with a 40×1 sensor resolution and 8 levels of sensitivity accurately traversed a curving IJ path over a landscape with a Gaussian blur of 1.0 (Fig 4a left) but failed early over a landscape with a blur of 10.0 (Fig 4a right). The interplay of landscape blur, sensor resolution, and sensor sensitivity on navigation success is shown in Fig 4b (note that RIDF but not TIDF was used in these trials). Blur factor and sensor sensitivity held more influence than sensor resolution (see the overlapping error bars for all resolutions, from 40×1 to 40×32). To simplify our results, we therefore averaged all six sensor resolutions to derive the 3D surface plot of Fig 4c, which is flattened to a 2D plot in Fig 4d. The results of all trials are available in S1 File.

**Fig 4.**
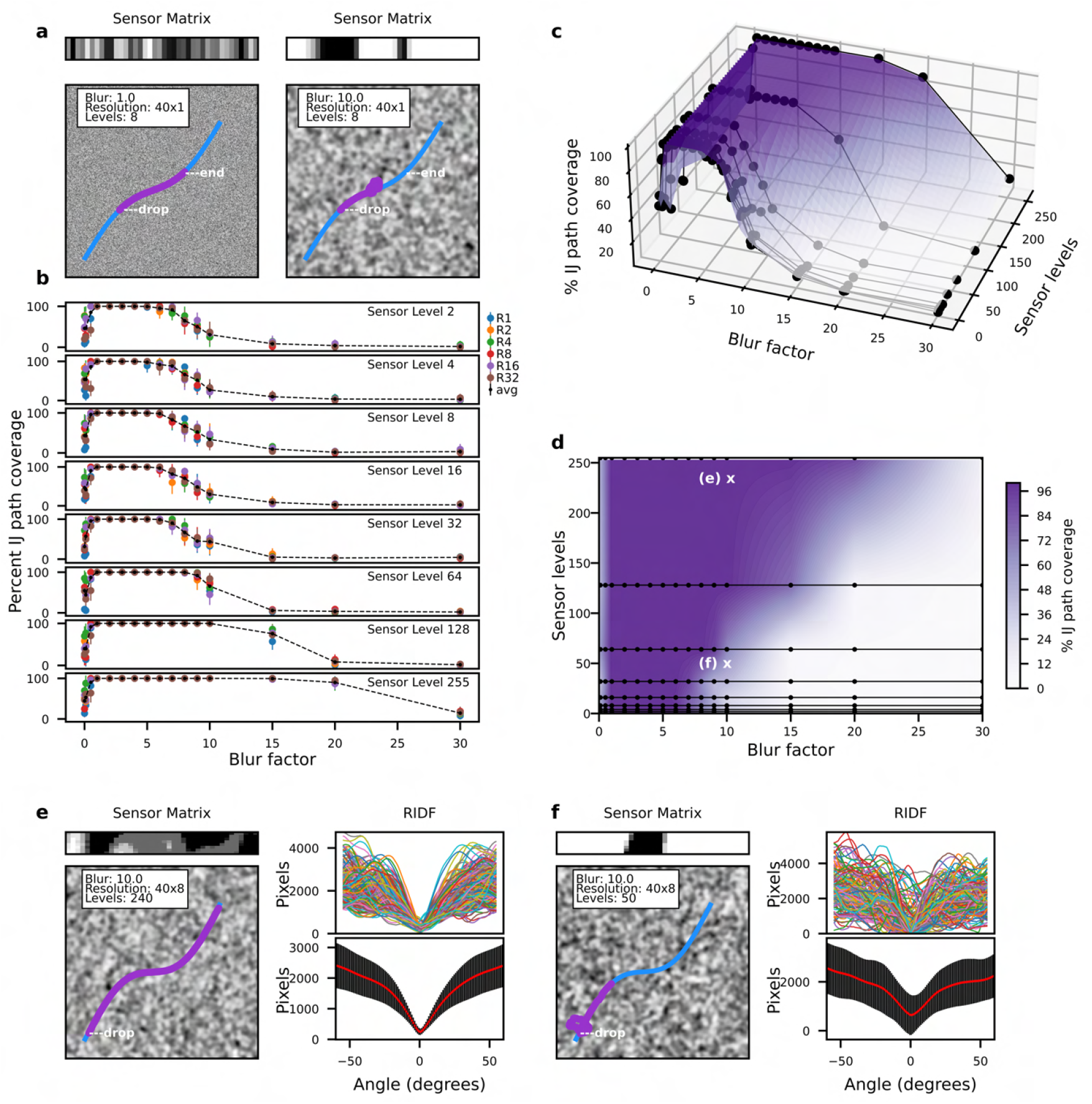
Sensor sensitivity and blur factor best predict navigation success. (a) Two SJ navigation examples (IJ curvature = 0.2; SJ start point = no offset from IJ; 100 SJ steps) with all parameters except blur factor held constant; left: blur=1.0, right: blur=10.0. (b) Percent path coverage (mean + CI) for all simulation runs plotted by blur factor (16 levels), sensor sensitivity level (8 grayscales), and sensor resolution (1, 2, 4, 8, 16, or 32 information units per tooth). Other parameters are the same as in (a). Each colored dot represents the average path coverage of 10 trials for each configuration. Black dots connected by dashed lines represent averages of sensor resolutions. (c) Surface plot of percent IJ path coverages for the averaged sensor resolutions (from b) against blur factor and sensor sensitivity level. (d) The same graph as (c) flattened to a two-dimensional plot. The two points marked with “x” have the same blur factor (10.0) but high (240 = ‘e’) and low (50 = ‘f’) sensor sensitivity levels. (e, f) Results of 250-step trials (step size = 4 pixels, resolution = 40×8, blur = 10, IJ curvature = 0.3, start point = 2 pixels offset from IJ) with sensor sensitivity of 240 1.or 50 (f). A sample sensor matrix with the relevant settings is shown above the landscapes. The upper plots on the right-hand side of e and f show superimposed RIDF plots (+60°) for the 250 SJ steps in each trial; lower plots show the average + SD of these plots.

To check the predictive power of various sensitivity/blur combinations, we compared navigation success using landscapes and sensors with characteristics taken from two points chosen from Fig 4d. One point was within the “high” part of the surface (~90% navigation success), with a blur of 10 and 240 sensor sensitivity levels. The other point was within the “lower” part of the surface (~24% navigation success) and had a blur of 10 and 50 sensor sensitivity levels. We tested navigation success using these combinations on curvier IJ paths than those depicted in Fig 4a and allowed the agents to complete the entire IJ path (250 total allowed steps). As predicted, the agent with parameters from the “high” part of the surface successfully navigated to the end of the IJ path (Fig 4e) while the agent with parameters from the “low” part of the surface failed soon after placement (Fig 4f).

### Testing NFLS across home range sized landscape

We tested whether NFLS can guide an agent along an IJ path similar to the size of the home range of a sand scorpion [18]. We found that an agent using both RIDF and TIDF successfully navigated a long (~2m) sinusoidal path (Fig 5, S2 Video). This figure also illustrates the random-walk behavior that typically happens after an agent on an SJ moves past its goal (the end of the IJ path). This behavior is a natural consequence of the scenes beyond the goal having no match in the agent’s memory. As such, the agent picks the best SAD from a set of poorly matched glimpses, with the meandering angle of each step being confined only by the width of the pre-set saccade angle.

**Fig 5.**
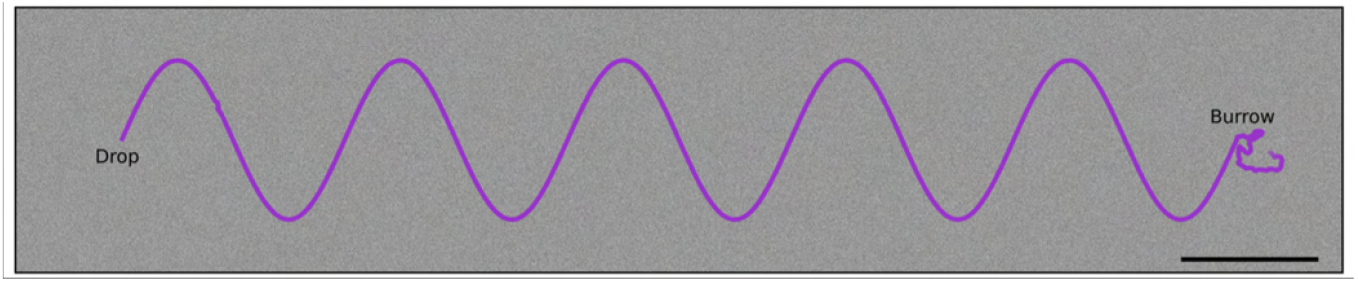
Navigation by familiarity works at scale of scorpion home range. An agent navigating by familiarity relative to a curved IJ path across a large simulated landscape (25,000×5,000 pixels). [blur = 2.0; sensitivity levels = 255; sensor resolution = 40×1; no. of steps = 4100; step length = 10 pixels; saccade width = +30°; lateral distance between TIDF samples = 4 pixels; scale bar = 10 cm]

### Assessing navigation success after displacement

Perhaps the most important consideration when assessing NFLS effectiveness is the lateral distance over which the IJ path influences the navigating agent. This measure considers the likelihood that an agent on an SJ using both TIDF and RIDF scene comparisons will find the IJ path when displaced laterally at various distances. A displaced agent navigating based solely on RIDF will tend to remain parallel to the IJ path. By adding translational information, the agent can choose the best SAD among three points (left, center, and right) and step forward from there. In this way the agent will gradually slide toward the bottom of the familiarity catchment trough after a few steps. In our tests we spaced the TIDF lateral samples at 2 pixels and used a blur/sensitivity combination that yielded essentially 100% navigation success when the agent was placed atop the IJ path in our previous tests (blur = 3.0, sensitivity = 255; resolution = 40×1). The results of these tests are shown in Fig 6. We used 100 unique landscapes with 25 start points spaced at 5 pixels on transects perpendicular and centered on the IJ path. The plot in Fig 6 shows the average success rate of the 100 points (+CI) at each distance from the IJ path using only a texture simulated landscape.

**Fig 6.**
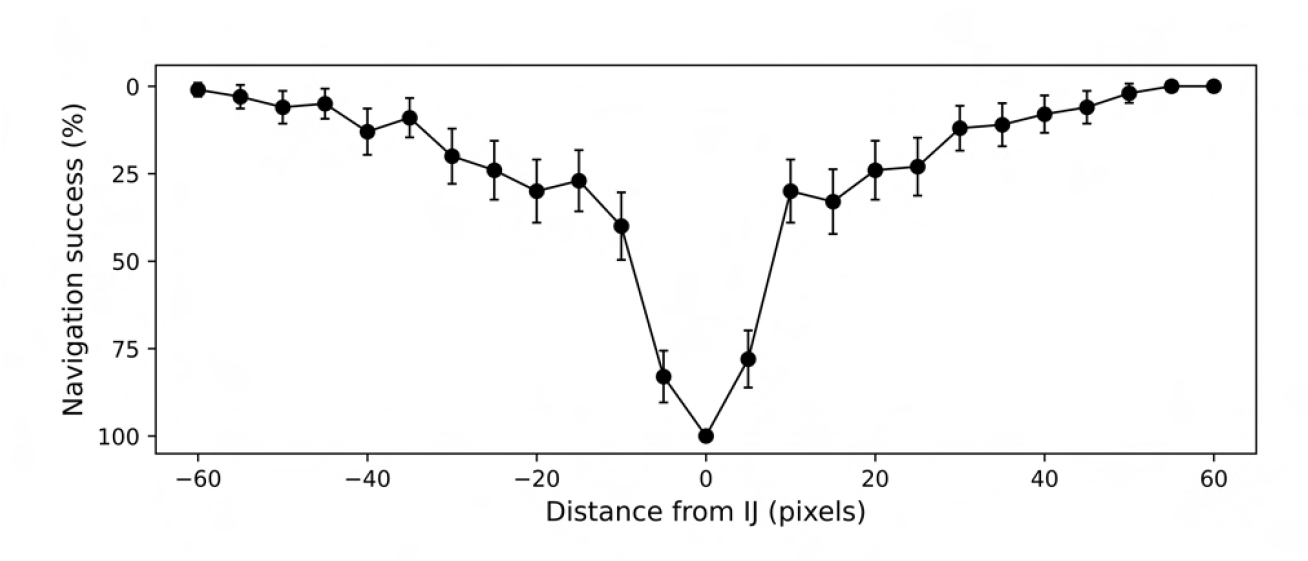
Navigation success after displacement. Results of navigation across texture only landscapes with agents started at various distances from the IJ path. Each trial used a different landscape (but with the same overall parameters: blur=3.0, sensor sensitivity levels=255, sensor resolution=40×1) and a straight IJ path (55 glimpses). The plot shows the average number of successful navigations (+CI) for each starting distance (n=100) from the IJ.

### Comparing single and multi-channel inputs on navigation success

Next, we assessed the effect of additional sensory channels on the ability of a displaced agent to find its way to the IJ path. In addition to the texture only landscape (used above), we ran 100 tests each of the texture landscape with independent additions of red, green, or blue (see materials and methods for details). As above, the plots in Fig 7a show the average (+CI) of the 100 points at each distance from the IJ path for each of the three conditions. Finally, we averaged the red, green, and blue results and compared the plot to the texture only navigation trials (Fig 7b). The width of this multi-sensory catchment trough at 50% success rate was 30.9 pixels (1.16 mm) while the texture only trough was only 15.0 pixels (0.56 mm). As such, the width of the catchment trough for the multisensory model was more than twice (30.9/15.0) as wide as the texture only (mechanosensory) trough. [Note: The results of all displacement tests (texture only and texture plus color) are available in S2 File.]

**Fig 7.**
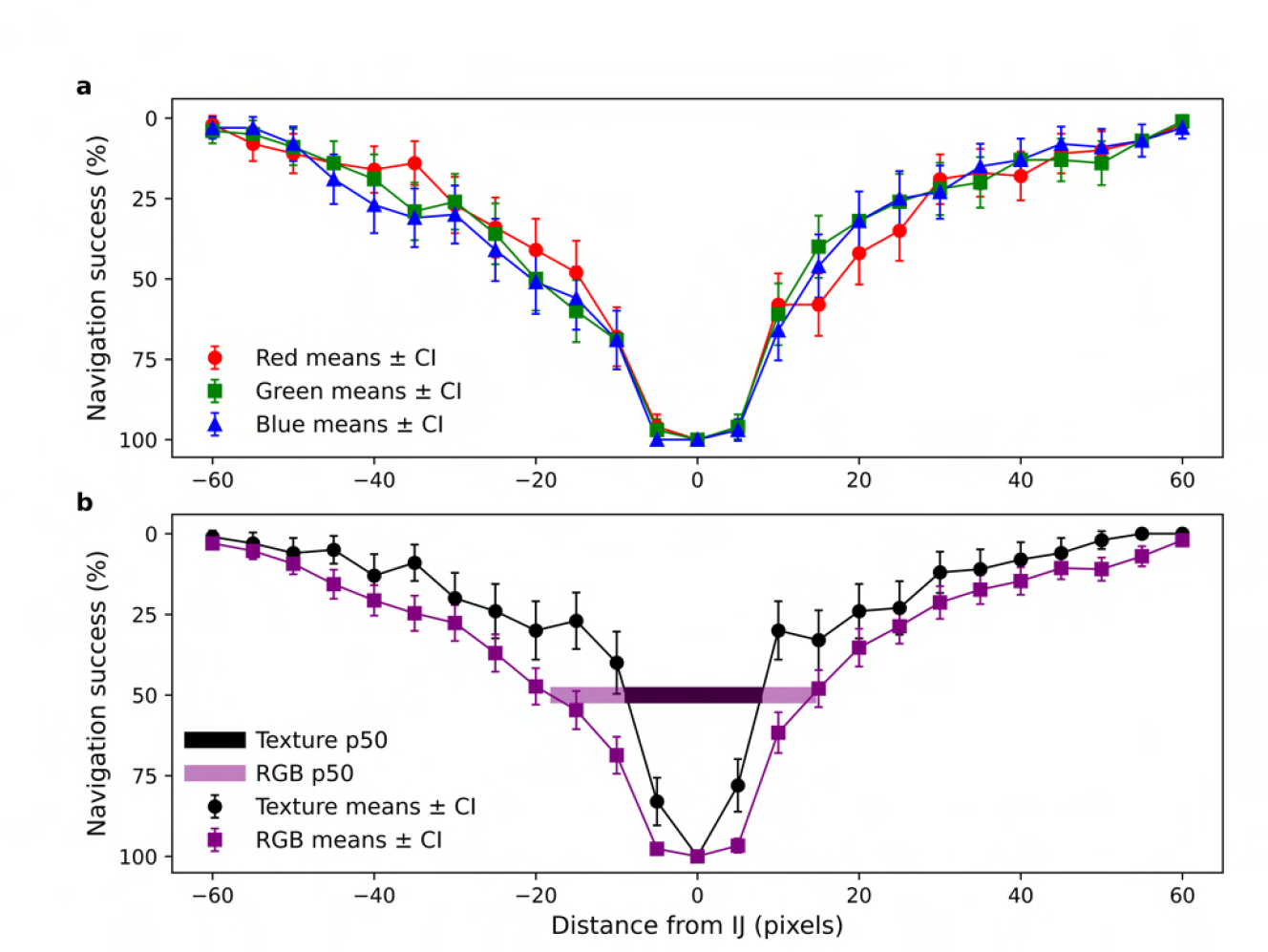
Multi-channel integration and catchment trough width. Results of navigation across texture only and texture plus colors (to simulate chemicals) landscapes with agents started at various distances from the IJ path. Each trial used a different landscape (but with the same overall parameters) and a straight IJ path (55 glimpses). The plots show the average number of successful navigations (+CI) for each starting distance (n=100) from the IJ. (a) Navigation results over 100 landscapes each for texture with added red (R), texture with added green (G) and texture with added blue (B) (see materials and methods for details of color generation and addition). (b) Navigation success of the averaged RGB trials compared to the texture only trials. The widths of the catchment troughs at 50% navigation success are shown (texture only: 15.0 pixels, RGB mean: 30.9 pixels); the relative increase in the catchment width of the multichannel vs single channel navigation is 2.06.

### Navigation amid landscape disruptions

Sand is vulnerable to displacement by wind, water, and the tracks of other animals. We therefore assessed the impact of landscape disruption on navigation success by overlaying an IJ path with various rectangular “patches” before an SJ began. Patches varied in both width and percent disruption of the underlying landscape. Fig 8 shows the results of 1430 trials where patch width and percent disruption were bracketed between 0 and 300 pixels (Fig 8a) and 0 and 100% (Fig 8b), respectively. Sample trials are shown in Fig 8c (the full set is provided in S3 Fig), and the results of these trials are summarized in the heat map of Fig 8d. All agents successfully completed the SJ when patches were less than 25 pixels wide or when the percent disruption was 20% or less. Beyond these values, navigation success fell in a graded manner as a product of both patch width and percent disruption.

**Fig 8.**
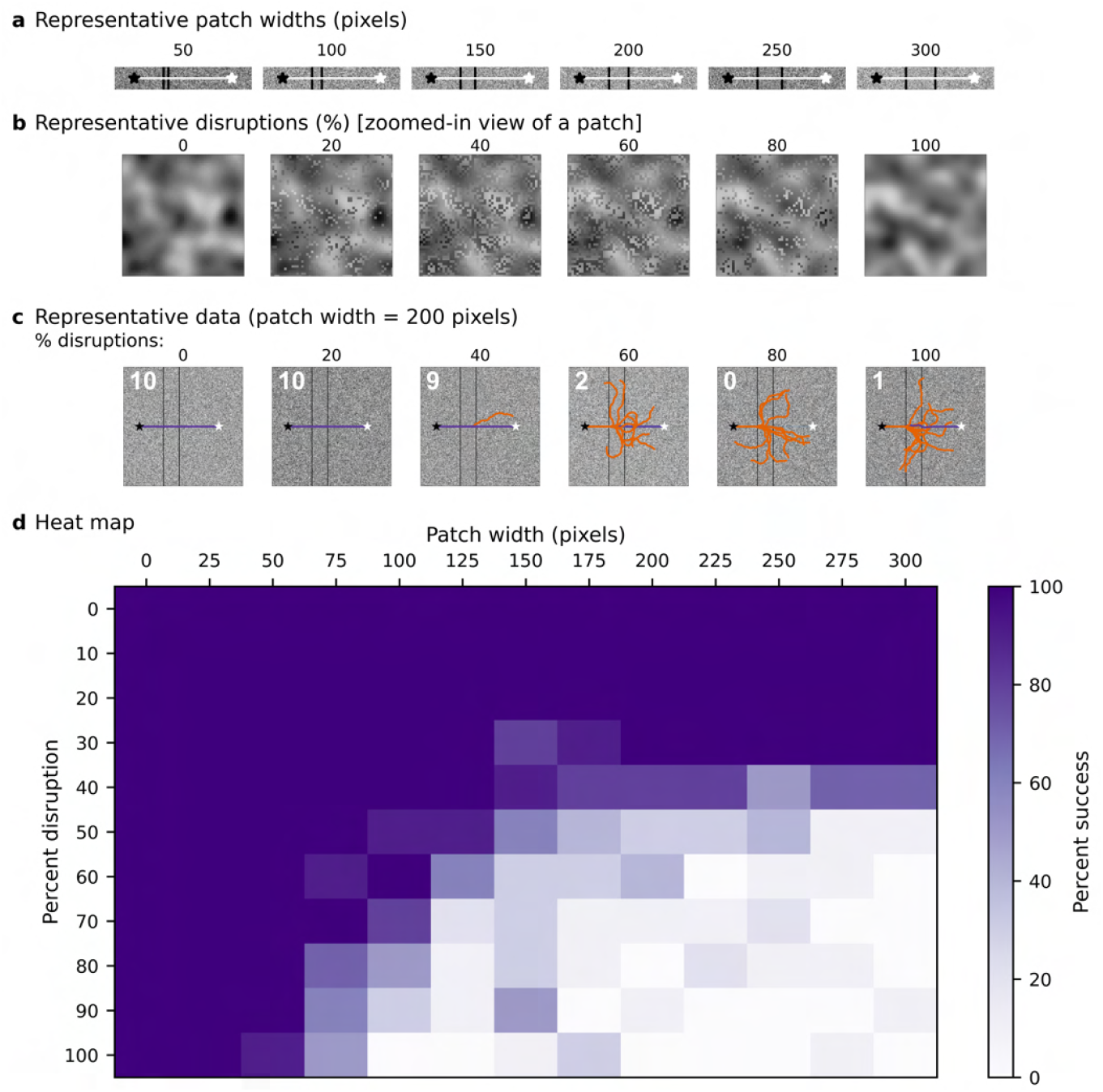
Testing navigation amid landscape disruptions. (a) Representative widths of disruptive rectangular patches across the IJ path (white line). (b) Expanded portions (50×50 pixels) of representative landscape disruptions from 0% (no disruption) to 100% (full disruption). (c) Examples of navigation success of agents on a straight IJ path with a patch width of 200 pixels and various levels of added landscape disruption (landscape blur = 3.0). The agent was dropped at the start of the IJ path (black stars), and each SJ was deemed successful if the agent moved within 10 pixels of the end of the IJ path (white stars). Successful SJ paths are color-coded purple, and unsuccessful paths are color-coded orange. Numbers in the upper left corner of each landscape indicate the number (out of 10) of successful trials. (d) The results of all trials are summarized in a heat map showing percent success for all patch widths and percent disruptions.

## Discussion

We have provided proof-of-concept evidence that navigation by familiarity as facilitated by scorpion pectines is both feasible and consistent with scorpion homing behavior. We have analyzed patterns in scorpion homing to self-dug burrows, and we have produced a computer simulation (NFLS) derived from physiological, morphological, and behavioral information about scorpions. We tested the model in a variety of navigational situations by systematically varying both landscape and sensory parameters to understand the interplay of these conditions on navigation performance. Most importantly, we have provided a framework for thinking about the dense matrices of peg sensilla on scorpion pectines and how they could form the basis of an elegant system for returning faithfully to an important site, such as shelter. This framework leads naturally to many additional, testable questions about the structure and function of not only the scorpion chemo-tactile pectinal system, but perhaps the function of overlooked sensory matrices on other animals.

### Do directed inbound routes suggest a transition from PI to familiarity navigation?

Our behavioral analyses showed that after executing a series of looping walks (putative learning walks) scorpions sometimes followed a consistent set of directed “inbound” routes to their burrow which were distinct from the dispersed “outbound” paths they took as they exited their burrow (see Fig 2). These observations, together with field [19] and laboratory studies, [20, 21] argue that the animals are using navigation by familiarity and not simply following so-called “breadcrumbs” in the form of footsteps or pheromones laid down as they move away from their burrow. Nonetheless, scorpions have been shown to use pheromones during mating [25, 28, 29] and are capable of mate-following [30], so it is hypothetically possible that some form of auto-chemosensation could be involved. For example, if animals leave deposits during short learning walks, it follows that chemical concentrations should be highest near the burrow. Such a gradient could be sensed by the pectines to guide burrow-directed behavior. While possible, such a system would be energetically costly since the animal must continuously produce and release significant amounts of signal molecules; further, the deposits would betray a scorpion’s existence and position to both prey and predators. Finally, dense matrices of peg sensilla, each with its population of 10+ sensory neurons, are overly complex and energetically expensive for the task of following a gradient by simple right/left comparisons, such as the chemical sampling of a snake’s forked tongue [59]. A more parsimonious interpretation is that scorpions use innate learning walks and path integration to scaffold home-directed glimpses of natural surface chemicals and textures for subsequent home navigation via familiarity.

Several additional points arise from the movement analysis of the animal in Fig 2 and S2 Fig. The animal made concentrated looping walks around its burrow during the first two hours following the burrow’s creation. Directed inbound routes began to form in the second hour and continued for the next five hours. The initial directed routes overlapped paths of the original looping walks but were straighter overall. The inbound paths led to the periphery of the mound at about 160° (on a 360° scale) while the actual burrow was about 60°. As such, the animals met the mound at a consistent spot, moved counterclockwise to the burrow, and returned to the arena wall from a variety of positions, mostly in the 0° to 90° range. The new directed paths overlapped previous paths, but with variation, which may be due to an imprecise path integrator. No matter the mechanism, this variation is useful from an NFLS standpoint. If each inbound journey adds to an animal’s collection of home-directed glimpses, a narrow pathway of familiar glimpses becomes a highway of recognition as the width of the familiarity catchment trough increases concomitantly [11, 60, 61]. Coupled with the animal’s forward momentum and limited saccade angles, corner-cutting on curvy paths should lead to straighter paths on subsequent journeys. Finally, it is possible that a trigger (neural or endocrine) could restrict new scene acquisitions to movements towards the burrow, thus coupling relevant scenes to the discharge of the animal’s path integration vector [9, 21].

### Limits to the pattern detection ability of the pectines

We hypothesize that the pectinal tooth (rather than the peg sensillum or entire pecten) is the unit of information in the scorpion CNS. Morphological studies show that projections from individual teeth coalesce in the pectinal neuropil and preserve the topographical order of the teeth relative to the ground [34, 35, 46]. Further, the spatial dimensions of the sand particles that make up the substrate of *P. utahensis* are more in line with the dimensions of the pectinal teeth compared to the peg sensilla [40, 47]. In the trials presented here, we found that agents with pectinal teeth sub-partitioned into regions (e.g., sensor resolution of 40×32) fared no better in our navigation tests compared to agents for which the tooth was the unit of information (i.e., sensor resolution of 40×1; Fig 4b). Finally, if the pectinal tooth is the unit of sensory information from the pecten to the CNS, then some mismatch might be tolerated by smoothing the data across the scores of pegs that adorn each tooth.

It is important to realize that a 40×1 resolution still yields enormous pattern recognition power. For example, if each tooth’s information output were simplified to a single bit, there would still be over a trillion matrix configurations. However, all evidence suggests the output from the tooth to the CNS is far from binary. The neural responses from individual peg sensilla are nuanced and capable of a complex repertoire of chemosensory responses based on chemical identity and concentration [20, 26, 38, 41]. In addition, the mechanosensory responses of the peg sensilla are also non-binary [20, 43]. By summing the tooth output based on the chemosensory and mechanosensory contributions of the hundreds of pegs on each tooth, the pattern recognition capacity of the pectines increases enormously. For example, if each tooth is capable of 100 output states instead of just two, pattern recognition capacity increases to 100^40^ or 1*10^80^! It is therefore not surprising that navigation success in our trials increased directly with sensor sensitivity from 2 to 255 (Fig 4c). Also, considering the graded chemo/mechano responses of the peg populations, the number of possible tooth states is likely to greatly exceed 255. This complexity suggests an interesting line of inquiry: Just how much information can be relayed by each tooth based on the summed response properties of tens to hundreds of pegs, and what are the limits of navigation success as tooth output increases accordingly?

One possible objection to our simulation is that biological systems (especially chemosensory systems [62] like the pectines) are vulnerable to sensory adaptation, whereas computer simulations are not. Sensory adaptation, in turn, could compromise accurate assessment and comparisons of chemical information. However, a local neural circuit in the pectines may mitigate this challenge. As alluded to before, peg sensilla contain numerous axo-axonic chemical synapses among the chemoreceptors just proximal to the sensory cell body layer in each tooth [33, 39, 50, 51]. These synapses appear to be part of a local negative feedback system that maintains peg neurons within a dynamic firing range, thereby overcoming adaptation [20, 52].

### Can scorpions navigating via NFLS overcome landscape disruptions?

A certain amount of substrate stability is required for a scorpion to navigate by NFLS, since the direction of each step is based on accurate matching of current glimpses with those stored in memory. Is the unconsolidated sand habitat of our main scorpion subject (*P. utahensis*) sufficiently stable? We began to address this question by assessing navigation success on disrupted landscapes (Fig 7). We found that even when a disrupted area was wide, the agent could successfully navigate the gap if the percent disruption was less than ~30%, akin to blowing or sprinkling a small amount of sand across an IJ path. In the Monahans Sand Hills area, where *P. utahensis* lives, the air and sand of the dune swales (where *P. utahensis* are most abundant; [18, 19]) are relatively stable at night but experience high heat and strong winds during the day (WillyWeather, National Weather Service). Photos of sand in the swales near scorpion burrows show that the footsteps of scorpions, beetles, and other nocturnal animals are still intact at sunrise but wiped clean by the afternoon. We suggest scorpions develop a new set of learning walks as they emerge each evening, somewhat akin to a spider that builds a new web daily. In addition, sand scorpions emerge to feed only a couple times a week on average [63, 64], and they tend to stay in their burrows during unfavorable weather [65, 66]. Perhaps they select emergence times to coincide with periods when conditions are most conducive to navigation by NFLS.

What if a strong wind blows a substantial amount of sand across the IJ path, or a slithering snake leaves a large swath of displaced sand in its wake? These are significant challenges with multiple possible solutions. First, adding multiple inbound paths (such as the directed inbound paths of Fig 2) would enable the animal to store more goal-directed glimpses in memory, which could overcome disruption of a portion of the landscape. Stored glimpses of other inbound paths generated by learning walks or longer inbound paths from previous excursions would also still be available. If a best match with memory is found with a scene that is part of an IJ path from another destination, the agent will be drawn into the alternate IJ catchment trough and guided to the burrow just the same [11].

Second, the path integrator vector should be available to move a confused animal in a homeward direction.

Third, meandering movements occurring after the agent moves past the goal (such as those beyond the end of the IJ in Fig 5) are reminiscent of the well-documented searching behavior in ants [67]. The meanders in our model are a natural consequence of poor matches between current glimpses and those in memory. Our algorithm could be adapted to execute expanding systematic loops as mismatch errors accumulate. Such a modification would increase the likelihood that an agent (or a scorpion) will encounter alternative familiar homebound paths. Preliminary tests (S4 Fig) show an agent with a fixed step length failing to regain the IJ (S3 Video) after it steps too far afield, whereas an agent whose step length varies indirectly with SAD value makes tight turns after stepping away from the IJ path and successfully regains the trail (S4 Video). It would be interesting to see if live scorpions use systematic searching when displaced from familiar territory.

Fourth, additional fault tolerance in NFLS navigation could be achieved through multisensory integration. As demonstrated in Fig 6, the addition of three sensory channels greatly improved navigation success. Although scorpion peg sensilla are sensitive to mechanical deflections [38, 43], the pegs are primarily chemosensors with neurons responsive to near-range stimulation by at least 20 organic chemicals [38]. We would like to assess the navigation performance in simulations equipped with chemical sensing capability approaching that of actual peg sensilla.

Fifth, vision could enhance familiarity navigation in scorpions. Early behavioral studies characterized scorpions as poorly visual [68, 69], based on results of trials conducted in unnatural light conditions. Dark-adapted scorpions, however, can readily detect light, even on moonless nights [70]. The paired median eyes, which sit atop the prosoma, should especially draw the notice of researchers interested in navigation [20]. They are composed of 500-600 retinular units [71] and have a 360° panoramic field of view [72]. Panoramic visual information from the median eyes may therefore be integrated with chemo-tactile information from the pectines to augment guidance along familiar paths, which would further compensate for disruptions to the sand surface tapestry.

Sixth, if all else fails, a wayward sand scorpion can always dig a new burrow [64].

### Other animals, other matrices

We think the power of the matrix has been overlooked in animal biology. QR code adoption was sluggish at first, but today the codes are ubiquitous [73]. We think the same could be true for understanding the matrices of sensilla that adorn the bodies of animals, especially those in Phylum Arthropoda. Nearly all arthropods – chelicerates and mandibulates alike – are covered with thousands of energetically expensive and dreadfully understudied sensilla. For example, why do the malleoli organs of solpugids have dense arrays of chemo-tactile sensilla [74, 75]? What about the exceptional antenniform first legs of amblypygids [20, 76, 77]? Could these arrays be involved in familiarity navigation in the animals’ native habitats? What about the arrays of sensilla on reproductive parts of many arthropod species? For example, could the arrays of mechanosensory sensilla on the thorax of female damselflies be involved in a system of template-matching of male mating parts and thus lie at the heart of sexual recognition and selection [78]? Are new worlds of sensory transduction and matrix processing awaiting discovery – for navigation, for object recognition, for communication, or for functions yet to be determined? We think so.

## Materials and methods

### Assessment of home-directed journeys

We revisited a set of all-night videos of sand scorpions (*Paruroctonus utahensis*) collected from the Walking Sands area of the Sevilleta National Wildlife Refuge in New Mexico, USA. The details of the laboratory setup are fully described in a previous paper [53]; in short, the animals were individually introduced into circular arenas (76.6 cm in diameter) lined with sand. Each arena contained a central mound of sand and we recorded the animals throughout the night with an overhead infrared video camera. Shortly after digging burrows in the sand mound, the scorpions made looping walks (putative learning walks) away from and back to the site of digging. In total, 23 animals were monitored, of which 18 produced usable recordings. All 18 showed evidence of learning walks.

In the current study, we wrote a MATLAB script to monitor the positions of these 18 animals beyond the times of the first learning walks and during subsequent evenings (for those that remained in the arena for multiple days). Of these 18 animals, 14 showed considerable movement throughout the arena. An additional MATLAB script was written for these 14 animals. First, a virtual circle superimposed on the image of the all-night tracking plot was used to delineate the mound region. The record was then queried for instances where the animal entered or exited the circle. “Inbound” paths were collected by excising the points of circle entry and their time stamps along with several seconds (typically 20) of the previous plot points. Similarly, “Outbound” paths were excised at the points of circle exit, along with several subsequent plot points (again typically 20). Of the 14 animals, 11 made multiple, overlapping movements from the arena wall to the central mound, suggesting directed navigational behavior. Plots of the movements of all 14 animals (representing 36 all-night records, since several animals were monitored for multiple nights), along with indications of directed walks, are shown in S1 Fig.

One animal displayed both learning walks and distinct, directed walks during its first evening of filming and its movements were selected for additional analyses. The inbound and outbound paths were further classified based on whether they extended to within a few centimeters of the arena wall. These wall-to-mound (W-M) and mound-to-wall (M-W) excursions suggested the animal was engaging in longer range foraging vs the looping patterns of the initial learning walks (see S2 Fig for in-depth analysis).

### Model of navigation by familiarity with a local sensor

We simulated scorpion navigation behavior using a computerized landscape and simulated pectines. We created our navigation simulations in Python (Spyder version: 5.4.3, conda) and our code is freely available at: [site provided on publication].

#### Landscape

Most tests used simplified, digitally generated, 1000×1000 pixel landscapes designed to visually simulate the texture of a sand scorpion’s habitat. Each pixel in these landscapes was randomly assigned a greyscale level ranging from 0 to 255. We also used a Gaussian blur function (*scipy*.*ndimage*.*gaussian-filter*) to create different levels of dispersion vs. coagulation among the pixels to generate landscapes that varied from fine to coarse in their particulate texture quality. Specifically, the blurring function applied a weighted average of surrounding pixels based on the Gaussian equation with larger factors causing greater coagulation. Later in the study, we added up to three color channels of information to the landscapes to simulate additional information carried by chemicals (described further below).

#### Sensor

We generated our simulated pectines based on the anatomy of the desert grassland scorpion, *Paruroctonus utahensis* (Williams, 1968), because substantial physiological and behavioral literature exits for this species [19, 20, 22, 23, 28, 40, 41, 43, 52, 53, 56]. These animals have sexual dimorphism in both the length of the pectines and the number of teeth per pecten. Males average 32.40 + 0.25 teeth per pecten (mean + SE; range 29 to 37, n=48, mode=33) and females average 19.88 + 0.16 teeth per pecten (mean + SE; range 17 to 22, n=42, mode=20) [79]. We conservatively modeled our agent’s sensor size based on the characteristics of female *P. utahensis* pectines. Further, female sand scorpions tend to be more faithful to their home burrows than are males, making them particularly useful subjects for the study of homing navigation (14). We examined the pectines of five adult female *P. utahensis* under a dissecting microscope and measured the width of the peg-containing surfaces from the inside of the most medial tooth to the outside of the most distal tooth; the average width was 3.00 + 0.08 mm and 3.10 + 0.06 mm (mean + SE) for the right and left pectines respectively. We estimated the anterior-posterior depth of the peg-containing tooth surfaces to be ~0.3 mm. We also measured the distance between the innermost tooth of the right pecten to the innermost tooth of the left pecten; the average was 3.45 + 0.15 mm (mean + SE) when the pectines are fully extended laterally from the body at 90°. The total distance from the most distal tooth of the right pecten to the most distal tooth of the left pecten is therefore ~9.55 mm (3.00 + 3.10 + 3.45). We also determined the number of teeth to be 20.60 + 0.93 and 20.60 + 0.81 (mean + SE) for the right and left pectines respectively. We therefore set the number of teeth on each simulated pecten at 20, or 40 for the combined pectines, which agrees with the statistical mode for this species from other studies [79].

We simplified our model pectines by removing the intervening gap and joining the right and left peg-containing tooth surfaces to form a rectangular bar that is 6.0 mm wide (rounding down from 6.1) and 0.3 mm deep. These dimensions created a 20:1 width to depth ratio (6.0/0.3=20). We used this ratio to produce model pectines of 160 pixels wide and 8 pixels deep (160/8=20). Each tooth of our model was therefore 8 pixels deep and 4 pixels wide (160 pixels / 40 teeth). This size allowed both for sufficient testing of navigational paths across our 1000×1000 pixel landscape and for generating several resolvable areas per tooth (see variables section for more details). The number of pixels per mm for our model can also be calculated as: 160 pixels / 6 mm = 26.67 pixels / mm. Each 1000×1000 pixel test landscape therefore had a physical size of 3.75 x 3.75 cm.

#### Variables used to assess the NFLS model

We tested navigation success and accuracy while varying the amount of information contained in the landscape (degree of blur), path characteristics (shape of the Initial Journey (IJ path) and the Subsequent Journey (SJ) initiation point), and sensor characteristics (sensor resolution, sensor sensitivity, saccade angles, and step length). These variables are summarized in Fig 9. In the tests designed to determine the combination of landscape, path, and sensor characteristics that best predict navigation success (Fig 4), we varied the parameters as follows. We held the IJ path curvature constant at 0.2, started the SJ at a consistent point atop the IJ, set the saccade to +60°, and fixed the step length to 4 pixels. We tested 16 levels of Gaussian blur in the landscape, 8 greyscales of sensor sensitivity, and 6 different sensor resolutions (40×1 to 40×32). We allowed each agent to complete 100 steps along the SJ and assessed the root-mean-square deviation (RMSD) and percent coverage of the SJ path relative to the IJ path (that is, the percent of the SJ path that fell within a step length [4 pixels] of the IJ path). We made 10 runs for each configuration (each with a new landscape) for a total of 7680 trials (16*8*6*10); data from all trials are available in the S1 File. Parameters for subsequent tests are described below.

**Fig 9.**
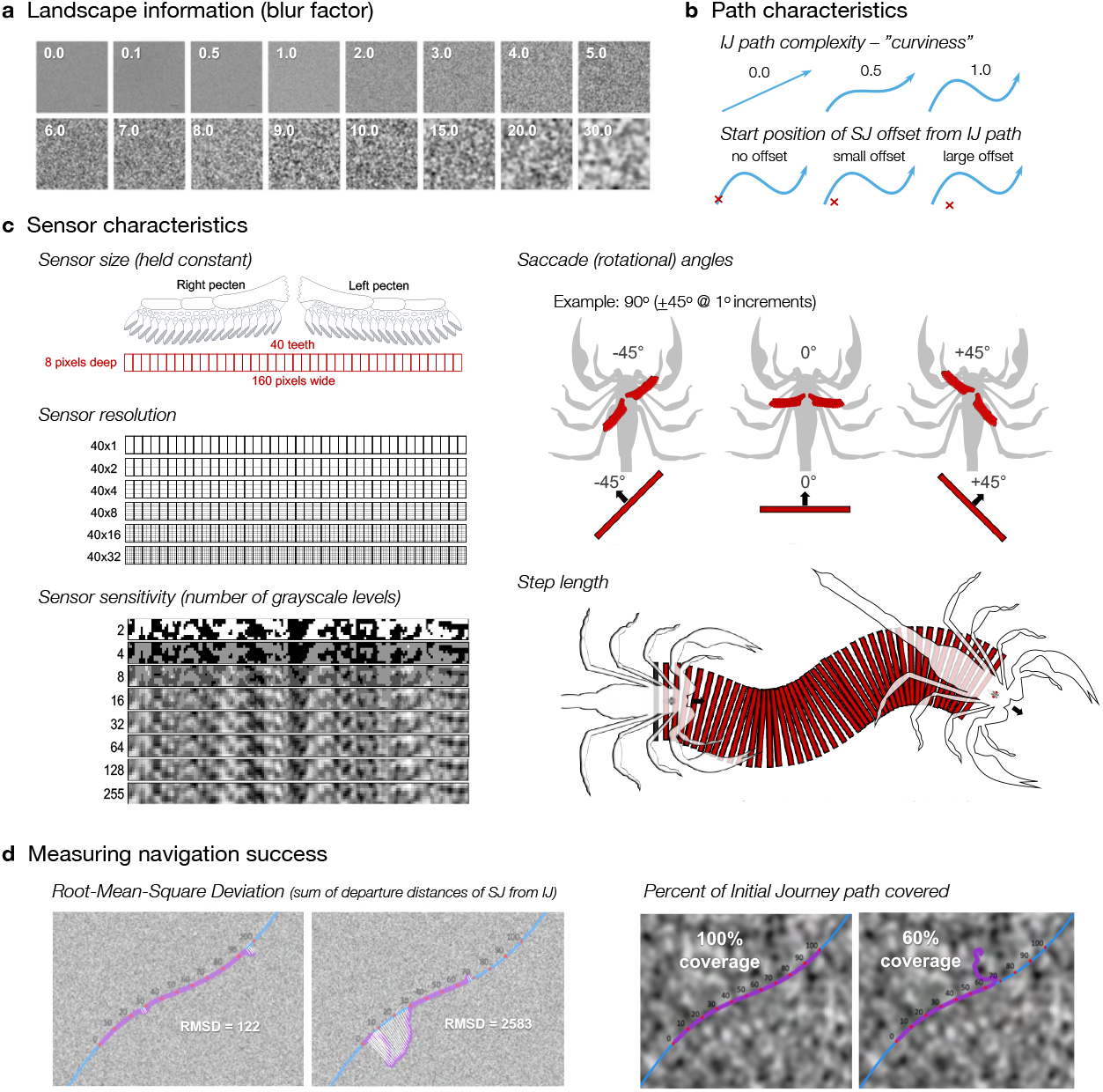
Variables used in testing the familiarity model. (a) Randomized greyscale landscapes were generated and passed through a Gaussian blur function with factors ranging from 0 (no blur) to 30 (heavy blur) at the 16 indicated levels. (b) The IJ path complexity varied from 0.0 (straight) to 1.0 (very curvy) and the start position of the SJ varied from directly atop the IJ to various distances lateral to the IJ. (c) We modeled the agent’s pectines after those of female *P. utahensis*, which average about 20 teeth on each pecten for a total of 40 teeth. We approximated the geometry of the peg-containing distal surfaces of the teeth to generate a rectangle of 8 pixels deep (~0.3 mm) and 160 pixels wide (~6.0 mm) for our simulated pectines (each tooth is 4 pixels wide). We left no space between the right and left pecten and simplified their rotational movements as if they were united as a bar. Sensor resolution varied from 40×1, where each tooth constituted a single unit of information, to 40×32, where each tooth was subdivided into 32 subunits of information. The 8 sensor sensitivity settings ranged from 2 to 255 greyscale levels. Saccade angles could be set to any right and left rotation value (in 1° increments; shown is a +45° saccade). Step length is the distance the agent moves forward between samplings (4 pixels for most trials in this study). (d) Navigation success was assessed by root-mean-square deviation (RMSD) and percent of the IJ path covered. In RMSD, the summed step-by-step distance of the SJ to the nearest points on the IJ points is calculated; the left panel shows a relatively successful SJ navigation with a low RMSD, whereas the SJ in the right panel deviated further from the IJ and has a much higher RMSD. The percent of the IJ’s length for which the SJ closely tracked the IJ was estimated at intervals of 10 percentage points. The panel at left shows a 100% coverage of the IJ by the SJ, whereas the panel at right shows only a 60% coverage.

#### Large-scale NFLS example

We challenged our simulated agent to navigate a distance akin to what an animal might experience in the field [18]. We used a very large texture-simulated landscape (25,000 pixels wide x 5,000 pixels tall), which is nearly a meter on its long edge (based on 258 pixels/cm). We used a single-channel sensor with sensitivity of 255 levels and resolution of 40×1. We created an undulating IJ path consisting of 20,000 glimpses. We started the agent 4 pixels off the beginning of the IJ path, set the step length to 10 pixels and the saccade to 60° degrees (+30° left/right), and allowed the agent to complete 4100 steps. For this simulation, we used both RIDF and TIDF, with the three TIDF points spaced at 4.0 pixels.

#### Assessing displacement from the IJ path

We also tested navigation success after starting the SJ systematically at various points offset from the IJ path. We first chose a landscape/sensor combination that reliably yielded successful navigations in previous tests: blur=3.0, sensor sensitivity levels=255, sensor resolution=40×1. The IJ was a short, straight path consisting of 55 captured scenes (spaced at 1 pixel). The SJ starting points formed a transect of 25 points centered on and extending in a line perpendicular to the IJ path. The transect crossed the IJ at 15 pixels from its beginning and the starting points were spaced at 5 pixels from each other. We allowed each agent to make 30 steps, at 2 pixels per step. An agent’s navigation was deemed successful if its SJ path moved within 2 pixels of the IJ path for three consecutive steps or if it came within 5 pixels of the end of the IJ path, whichever came first. For these simulations, we used both RIDF and TIDF, with the three TIDF points spaced at 2.0 pixels. We then plotted the average number of successful navigations (+ CI) for each starting distance from the IJ. This process yielded an average “catchment trough,” which helped us determine the influence of the IJ path scenes at various distances from the IJ.

#### Multi-channel integration

We assessed the effect of multisensory integration on the depth and width of each catchment trough by adding additional channels of information. Specifically, we added three color channels (red, green, blue) to simulate information carried by contact chemoreceptors that would be detecting three different types of chemicals. We began with the same 100 texture-only (greyscale) landscapes used in the displacement experiments. To integrate color, we also created 100 new landscapes consisting of randomly placed, overlapping, randomly sized circles (between 20 – 50 pixels in radius). Each circle was assigned a color by randomly selecting a value of 0 to 255 for each of the three components of an RGB color triplet. We applied a Gaussian blur (blur factor = 71) to each color landscape to simulate the dispersion of chemical stimuli. Finally, we separated each chemical-simulated landscape into three color-isolated channels (R, G, and B). We paired each chemical-simulated landscape with a texture-only landscape, and we separately superimposed each color on the corresponding simulated texture-only landscape. As before, we created transects of 25 SJ initiation points and plotted the average number of successful navigations (+ CI) for each starting distance from the IJ for each of the color-added landscapes, as well as the average of the three colors. In all, we used 100 trials for our tests for a total of 10000 navigation runs (4 landscape conditions * 25 initiation points * 100 trials). All trials and their accompanying runs are archived in S2 File with successful SJ paths colored purple and unsuccessful paths colored orange.

#### Testing navigation amid landscape disruptions

We generated a set of landscapes with various levels of disruption across the IJ path. Specifically, we first generated 2000×2000 pixel, single channel (i.e., texture-simulated) landscapes with a Gaussian blur of 3.0. Across the middle of each landscape, we made a straight IJ path consisting of 1000 glimpses, spaced 1 pixel apart. Disruptions were then imposed on the landscapes by placing rectangular patches of varying widths perpendicular to the IJ path, beginning 300 pixels from the start of the IJ. These patches were generated anew by the same mechanism that produced the 3.0 Gaussian blur of the original landscape. The patches spanned the 2000-pixel vertical dimension of the original landscape and varied in width from 0 pixels (no disruption) to 300 pixels (large disruption) in 25-pixel increments, for a total of 13 different patch widths.

The patches were further modified to vary the percentage of the patch’s landscape that was substituted for the original landscape. For example, a patch with 40% disruption had randomly selected pixels substituted for 40% of the original landscape pixels and retained 60% of the original landscape pixels. We varied the percent disruption from 0% (no disruption) to 100% (full disruption) at 10% increments for a total of 11 different percent disruptions.

We assessed navigation success by dropping agents with sensor resolutions of 40×1 and 255 levels of sensitivity at the beginning of the IJ path. The saccade angle was set at 60° (+30° left/right) and both RIDF and TIDF were used (translation glimpses spaced at 1 pixel left and right). Each trial consisted of 50 completed steps at 20 pixels per step, and 10 trials were run for each landscape, with a new landscape (and patch) being generated between trials. A trial was deemed successful if the SJ took the agent within 10 pixels of the end of the IJ path; our dependent variable was the number of successful trials. In all, this portion of the study consisted of 143 landscapes (13*11) and 1430 total trials (these trials are shown in S3 Fig).

## Supporting information

NOTE: Repositories of our Python code will be provided upon publication. All additional data are available in the main text or the supplementary materials.

**S1 Fig**. Examples of learning walks and directed movements in all-night plots of scorpion activity.

**S2 Fig**. In depth movement analysis of animal shown in Fig 2.

**S3 Fig**. Simulation trials supporting landscape disruption tests of Fig 8.

**S4 Fig**. Long range navigation simulations using agents with fixed vs. dynamic step lengths.

**S1 Video**. Animated simulation of simplified NFLS model shown in figure 3.

**S2 Video**. Animated simulation of the large-scale navigation example shown in figure 5.

**S3 Video**. Animation of the example depicted in Supplemental Figure 4a.

**S4 Video**. Animation of the example depicted in Supplemental Figure 4b.

**S1 File**. Replication data supporting tests of landscape and sensor variables depicted in Fig 4.

**S2 File**. Replication data supporting displacement and multi-channel trials in Fig 6 and Fig 7.

## Acknowledgments

We thank Albert Musaelian for inspiration concerning the Python implementation of the NFLS algorithm and Paul Graham for critical review of early drafts of this manuscript. We also thank George Martin for engineering assistance, the staff of OU’s Laboratory Animal Resources for assistance with animal care, the OU School of Biological Sciences for facilities support, and OU Presidential Teaching Fellow in Honors funds to DDG for research materials. Finally, we thank the UNM Sevilleta Field Station and personnel for lodging and research support and the Sevilleta LTER for access to field sites.

