## Supplemental Figure 3 for "Scorpion navigation by chemo-textural familiarity: modeling the interplay between sensory and landscape parameters"

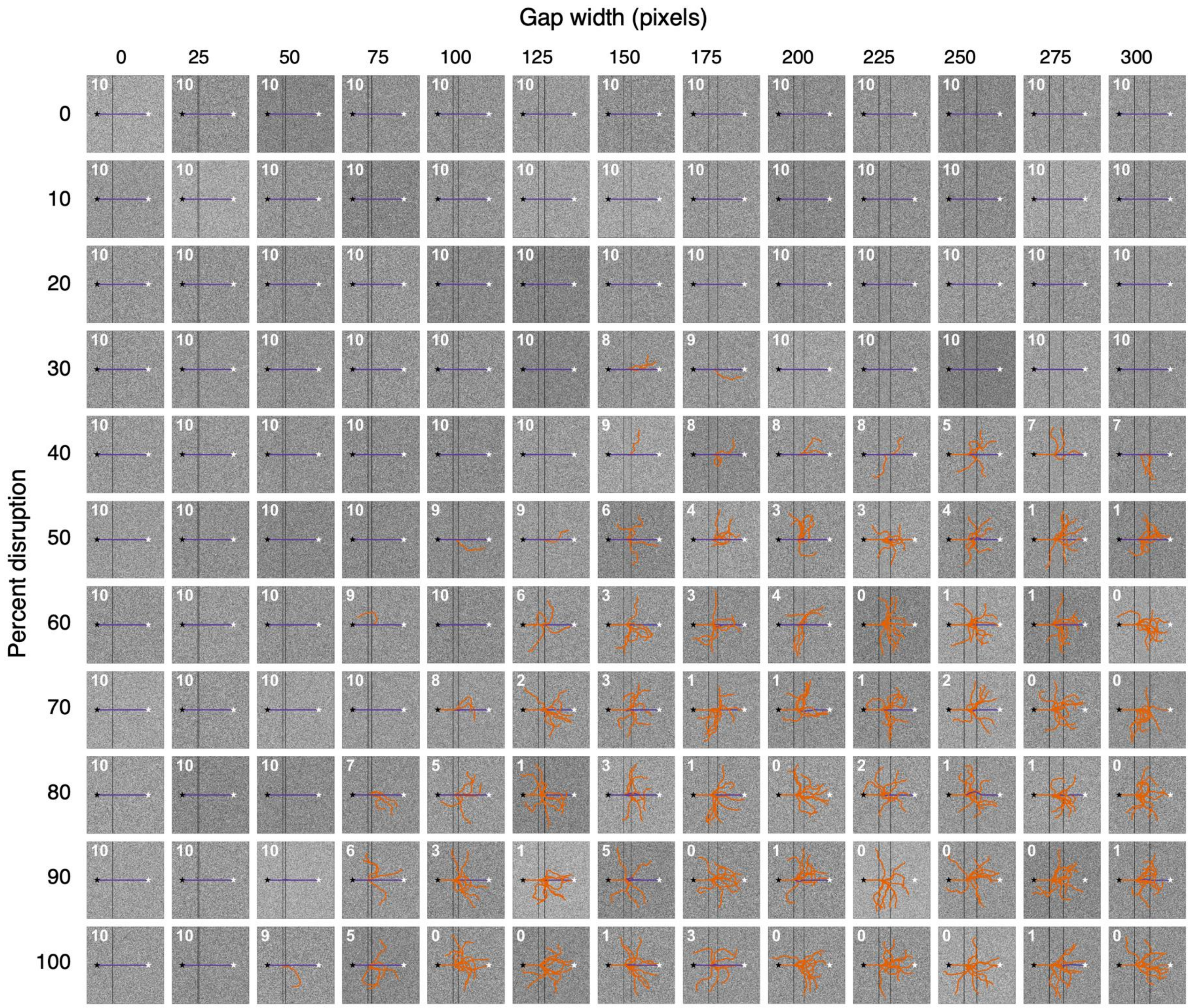


**SF3 Supporting data for Fig 8.** The navigation success of an agent using both RIDF and TIDF is assessed on a straight IJ path across texture-simulated landscapes (2000x2000 pixels; blur = 3.0) with various levels of added environmental disruption. The agent was dropped at the start of the IJ path (black stars) and deemed successful if it moved within 10 pixels of the end of the IJ path (white stars). Disruptions consisted of altered patches of landscape arranged perpendicular to the IJ path ranging from 0 to 300 pixels in gap width in steps of 25 pixels. Each patch was further modified based on percent disruption from 0% to 100% at 10% intervals. The agent's sensitivity, resolution, and saccade angle were held constant at 255, 40x1, and 60$^{o}$ respectively; the distance between TIDF samples was set at 1 pixel left and right. Each agent ran 10 trials for each disruption with each trial consisting of 50 steps at a step length of 20 pixels. A new landscape was generated for every trial. Successful SJ paths are color-coded purple while unsuccessful paths are color-coded orange. Numbers in the upper left corner of each landscape indicate the number (out of 10) of successful trials.
