## Supplemental Figure 4 for "Scorpion navigation by chemo-textural familiarity: modeling the interplay between sensory and landscape parameters"

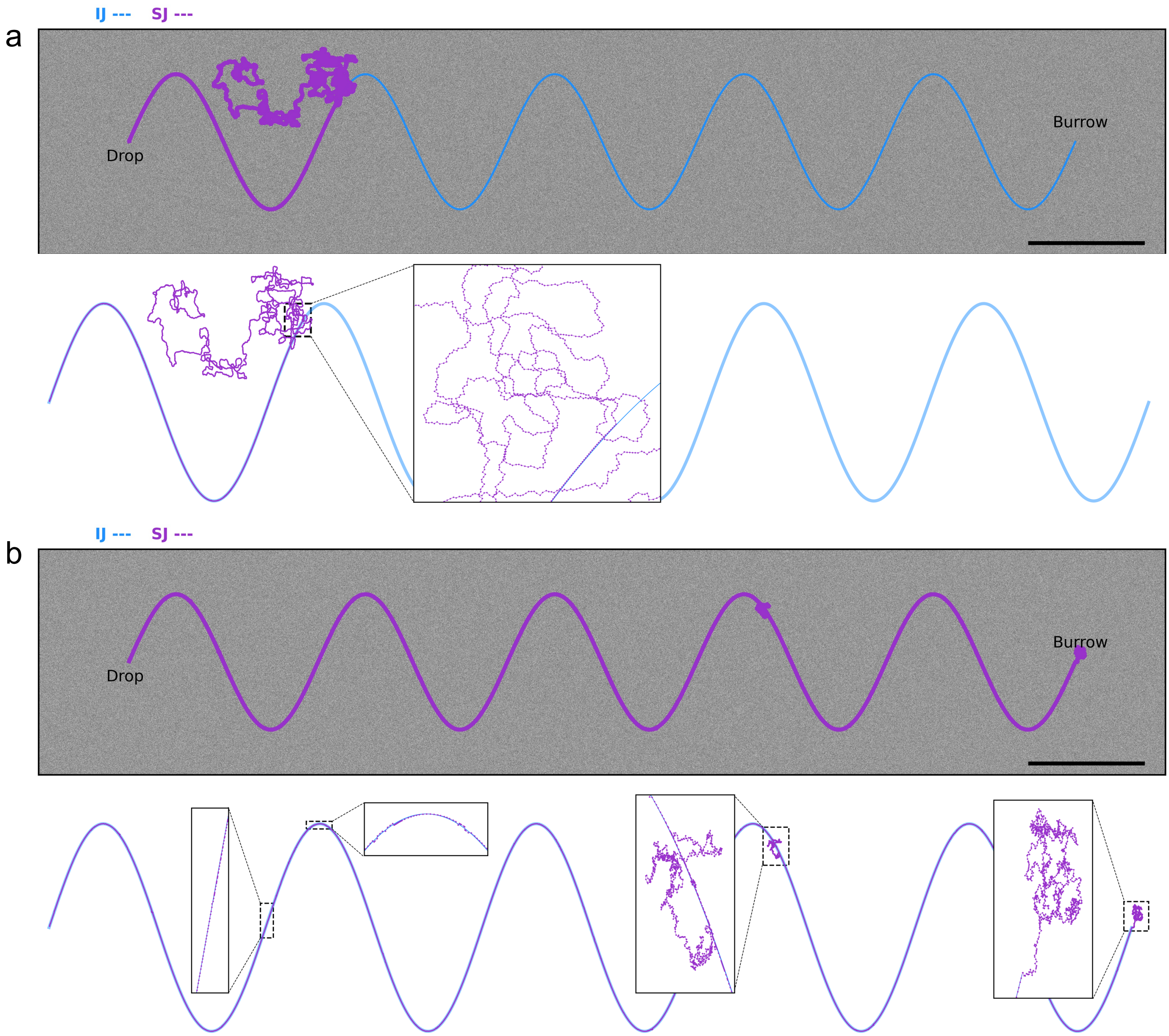


**SF4 Comparison of agents with fixed vs. dynamic step length relative to SAD value.** (a) An example of a navigation failure of an agent with a fixed step length (5 pixels) navigating relative to a curved IJ path across a 25,000x5,000 pixels texture-simulated landscape. [blur = 2.0; sensitivity levels = 255; sensor resolution = 40x1; no. of steps = 8000; step length = 5 pixels; saccade width = 60º; lateral distance between TIDF samples = 4 pixels; scale bar = 10 cm]. (b) An example of a successful navigation by an agent with a dynamic step length that varied indirectly with SAD value. Also, the saccade width of this agent was increased to 120º and the lateral distance between TIDF varied directly with SAD. [blur = 2.0; sensitivity levels = 255; sensor resolution = 40x1; no. of steps = 8000; scale bar = 10 cm].
